# Automated identification of chicken distress vocalisations using deep learning models

**DOI:** 10.1101/2021.12.15.472774

**Authors:** Axiu Mao, Claire Giraudet, Kai Liu, Inês De Almeida Nolasco, Zhiqin Xie, Zhixun Xie, Yue Gao, James Theobald, Devaki Bhatta, Rebecca Stewart, Alan G. McElligott

**Author notes:** **Corresponding authors:** KL;, AGM.

## Abstract

The annual global production of chickens exceeds 25 billion birds, and they are often housed in very large groups, numbering thousands. Distress calling triggered by various sources of stress has been suggested as an “iceberg indicator” of chicken welfare. However, to date, the identification of distress calls largely relies on manual annotations, which is very labour-intensive and time-consuming. Thus, a novel light-VGG11 was developed to automatically identify chicken distress calls using recordings (3,363 distress calls and 1,973 natural barn sounds) collected on intensive chicken farms. The light-VGG11 was modified from VGG11 with a significantly smaller size in parameters (9.3 million *vs* 128 million) and 55.88% faster detection speed while displaying comparable performance, *i*.*e*., precision (94.58%), recall (94.89%), F1-score (94.73%), and accuracy (95.07%), therefore more useful for model deployment in practice. To further improve the light-VGG11’s performance, we investigated the impacts of different data augmentation techniques (*i*.*e*., time masking, frequency masking, mixed spectrograms of the same class, and Gaussian noise) and found that they could improve distress calls detection by up to 1.52%. In terms of precision livestock farming, our research opens new opportunities for developing technologies used to monitor the output of distress calls in large, commercial chicken flocks.

## 1. INTRODUCTION

Vocalisations can be used to infer whether animals are experiencing positive or negative states [1,2]. Chickens (*Gallus gallus domesticus*) have a variety of vocalisations associated with different states, including pleasure and distress [3,4]. Distress vocalisations made by young chickens are repetitive, high-energy, and relatively loud calls [5]. Importantly, the output of distress calls from commercial broiler chicken flocks (*e*.*g*., approximately 25,000 birds together in one location) can be used to predict growth rates and mortality levels during the production cycle [5]. However, to date, the process of assessing the number of distress calls produced in large-scale recordings largely relies on manual annotations, which is labour-intensive, time-consuming, and prone to subjective judgments of individuals. Thus, it is essential to develop new automated, objective, and cost-effective methods for identifying and quantifying distress vocalisations, against a background of other vocalisations and noises that are usually contained in the audio recordings.

Despite research highlighting the potential for automated monitoring of vocalisations as a means to assess and monitor animal welfare states [6], progress has been slow. In chickens, most methods have focused on detecting issues associated with respiratory diseases or measuring growth [7–10]. However, due to the links between emotional states and types of vocalisations and recent advances in machine learning applied to audio data [3,11–15], we hypothesised that automated detection of chicken distress calls would be feasible.

The use of bioacoustic methods as non-invasive techniques for monitoring health and welfare is becoming more widespread [12,13]. For distinguishing healthy birds from those with infectious bronchitis, a support vector machine (SVM) was used to reach 97.85% accuracy based on their audio recordings [10]. Sneezing is a clinical sign of many respiratory diseases, and the monitoring of which was achieved through a linear discriminant analysis (LDA) using chicken sound signals at a group level [16]. Du *et al*. [17] were the first to show a significant correlation between specific vocalisations (alarm and squawk call) and thermal comfort indices (temperature-humidity index), using a hen vocalisation detection algorithm based on SVM. However, manual feature extraction in machine learning relies on expert domains and easily causes feature engineering issues [18]. Furthermore, feature extraction and classification separation makes the process unsuitable for online and real-time large data analysis [19].

Recently, deep learning classification models have been highly accurate and reliable in bioacoustics [15,20–22]. In particular, convolutional neural networks (CNNs) have been used predominantly to process image data and other image-like data, such as Mel spectrograms, the most common input for deep learning models [12]. A spectrogram can visually represent signal strength over time at different frequencies present in a specific waveform [23]. In practice, CNN-based models, including some existing CNN architectures (*e*.*g*., AlexNet, VGG-16, and ResNet50), have been used to recognise a wide variety of vocalisations (*e*.*g*., pecking activity of group-housed turkeys, alarm calls of laying hens, and cough sounds of cattle) using audio spectrograms [21,24,25].

In real-world scenarios, the available data for training is rather limited, either due to the enormous manual efforts required to collect and annotate data or since it is difficult to acquire large amounts of data in some cases [25]. Data augmentation as an elegant solution has been beneficial for expanding dataset sizes [26]. It artificially enlarges the data representativeness through label-preserving transformations to simulate real-world scenarios with the presence of many different kinds of noise [27]. The data augmentation techniques commonly used in image data comprise random cropping, rotation, flip, and scaling [26]. In bioacoustics, when a time-frequency sound signal representation obtains images, the data augmentation techniques must be specifically devised for this context [27]. For bird audio detection using CNN-based models, time shift, frequency shift, and time reversing were randomly applied on the spectrograms during training to reduce overfitting [28]. Recently, an augmentation scheme named SpecAugment, which acts exclusively on the audio spectrogram, has been validated to enhance the performance of end-to-end networks on public speech datasets [29,30].

In this study, we exploited deep learning models to classify audio recordings of large groups housed indoors on commercial chicken farms, to investigate the potential of deep learning for automatically identifying chicken distress calls. To automatically classify distress calls and other natural background sounds recorded, we developed a novel light-VGG11 model modified from VGG11 with a significantly smaller size in parameters (9.3 million *vs* 128 million). In addition, to improve the generalisation capability of the models, data augmentation was explored as a means to increase the diversity of the limited training dataset. To the best of our knowledge, this is the first time deep learning methods were utilised in chicken distress calls identification based on audio recordings.

## 2. METHODS

### 2.1. Data Collection

Recordings were collected in production facilities owned by Lengfeng Poultry Ltd., in Guangxi province, People’s Republic of China, between November and December 2017 and in November 2018. Chickens (mix of Chinese “spotted” and “three-yellow” breeds) were kept in stacked cages (three cages per stack, with 13-20 individuals per cage), with approximately 2,000 to 2,500 birds per house, as shown in Figure 1. The microphone was positioned approximately two meters up from the floor, and attached to barred windows. This placement was chosen to ensure that the recording devices did not interfere with the farm staff cleaning and maintaining the barns. All recordings were sampled at a 22.05 kHz with a 16-bit resolution throughout the natural lifecycle (0 – 35 days) of chickens using a Zoom H4n Pro Portable Recorder (Zoom Corporation, Tokyo, Japan).

**Figure 1.**
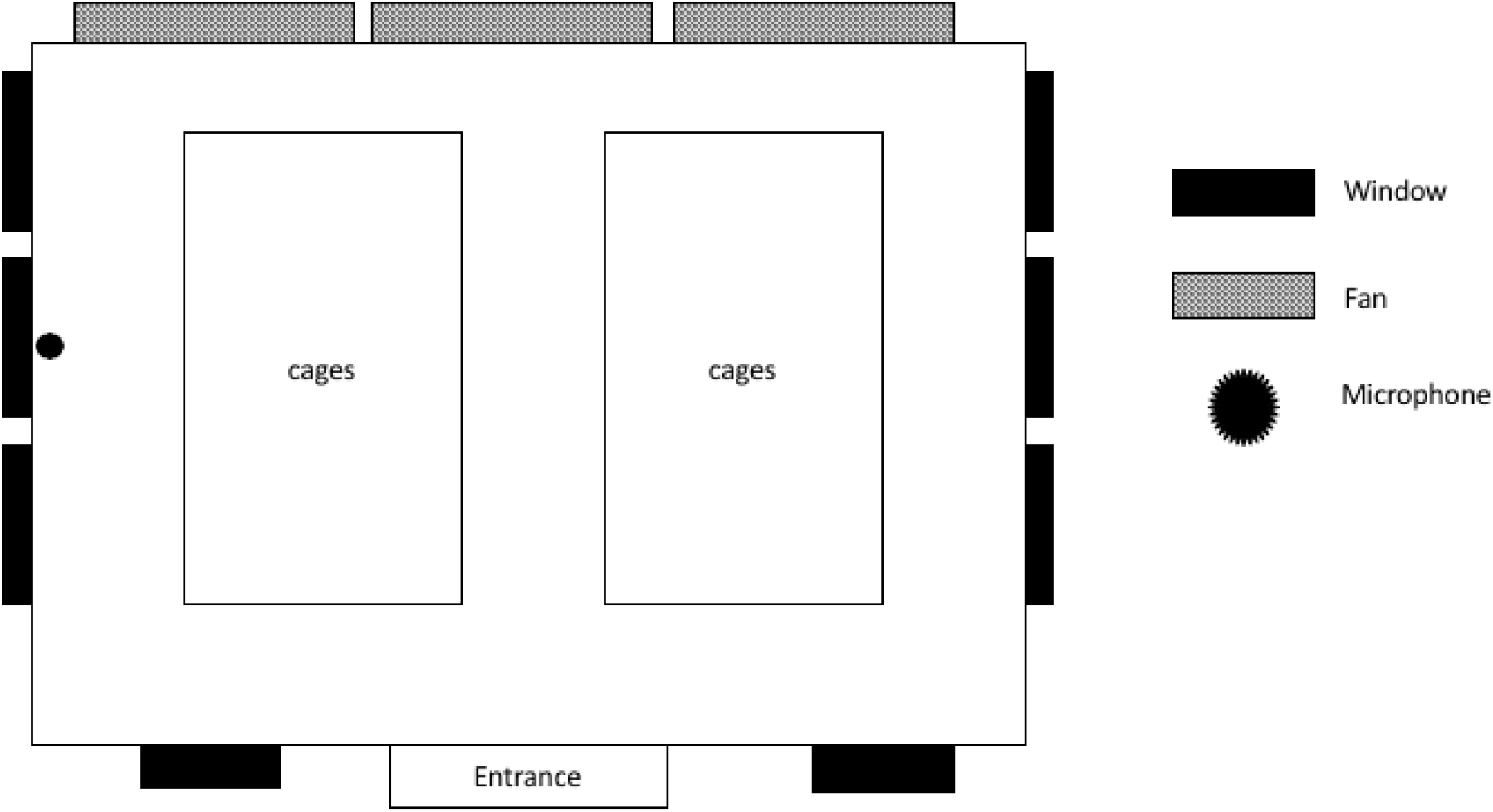
The layout of the chicken house and microphone placement (84 m length, 11.3 m width, and m height).

### 2.2. Data Annotation

Data were annotated using the Sonic Visualiser software (Sonic Visualiser Version 4.3, Centre for Digital Music, Queen Mary University of London, UK)[31].The audio taxonomy consisted of distress calls, work sounds, and natural barn sounds. Work sounds were defined as any noises that the farm staff made while cleaning the barn or carrying out routine husbandry for the birds. Natural barn sounds were the natural sounds of the barn when there were neither distress calls nor work sounds inside the barn. Calls such as pleasure chirps and fear trills were not annotated due to their relatively low amplitude compared to distress calls [32]. Contact calls were not included because they are produced nearly continually by the animals and are not indicative of the welfare aspect [32]. In particular, distress calls and natural barn sounds were the sounds of interest, and their examples of data annotation are shown in Figure 2. The labels and timestamps of these two types sound of interest were then exported to CSV files for further processing.

**Figure 2.**
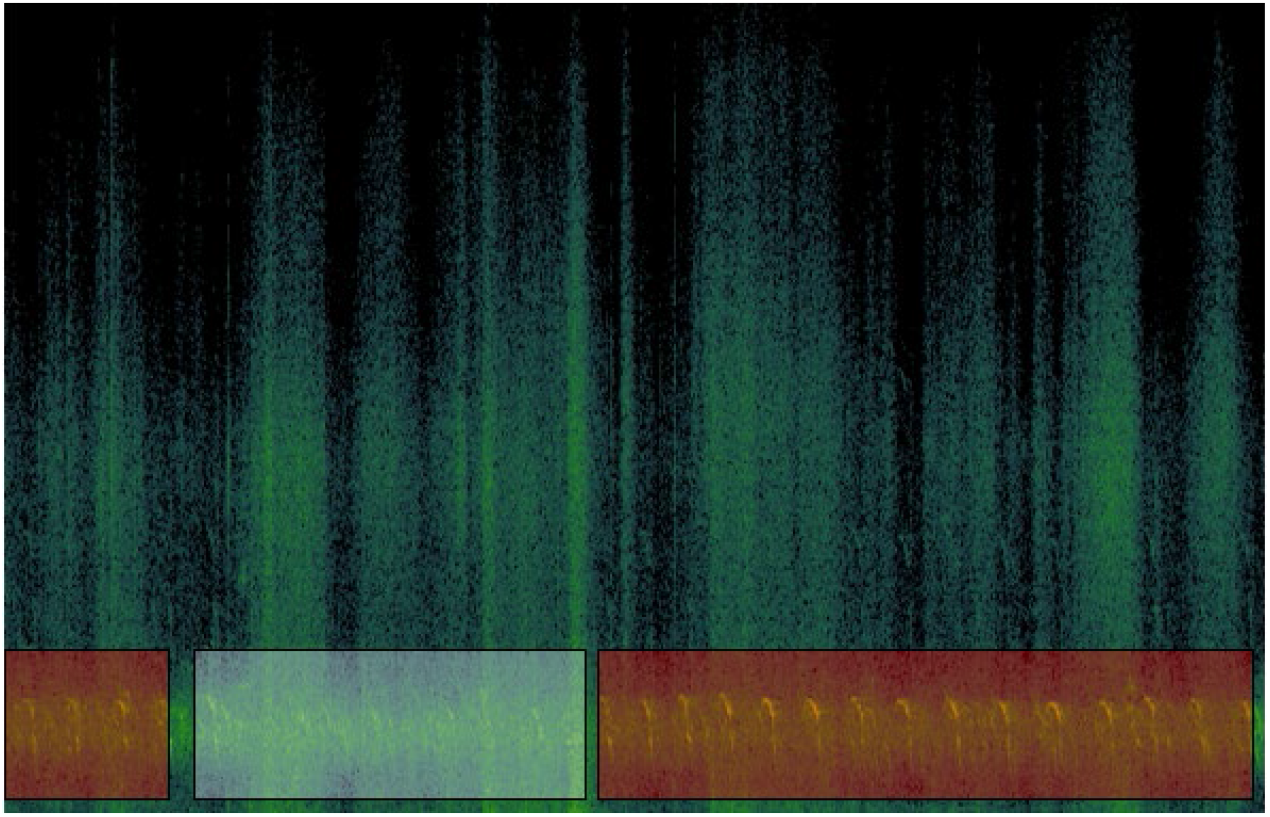
An example of how the data were annotated. The squares in red highlight the distress calls, while the square in white shows what would be termed ‘natural barn sounds’ (the absence of work sounds and distress calls).

### 2.3. Automatic Classification

We developed an audio classification method based on deep learning, *i*.*e*., light-VGG11 (see  section 2.3.2), and a bioacoustic dataset (Figure 3). First, recordings in the audio files were chunked into non-overlapping segments of one second to produce a set of 5,336 samples, which consist of 3,363 distress calls and 1,973 natural barn sounds (see Data Accessibility Section). The segments less than one second long were excluded from the dataset. These raw audio segments were then transformed to log-Mel spectrograms, then image-wise normalised to the range [0,1]. After that, we split these normalised spectrograms into training, validation, and test sets based on a five-fold cross-validation technique that helps avoid overfitting and promotes the model’s robustness. The training set and validation set were utilised to learn the model, and the test set was used to test the model. The average of the five test results was used to evaluate the model’s performance.

**Figure 3.**
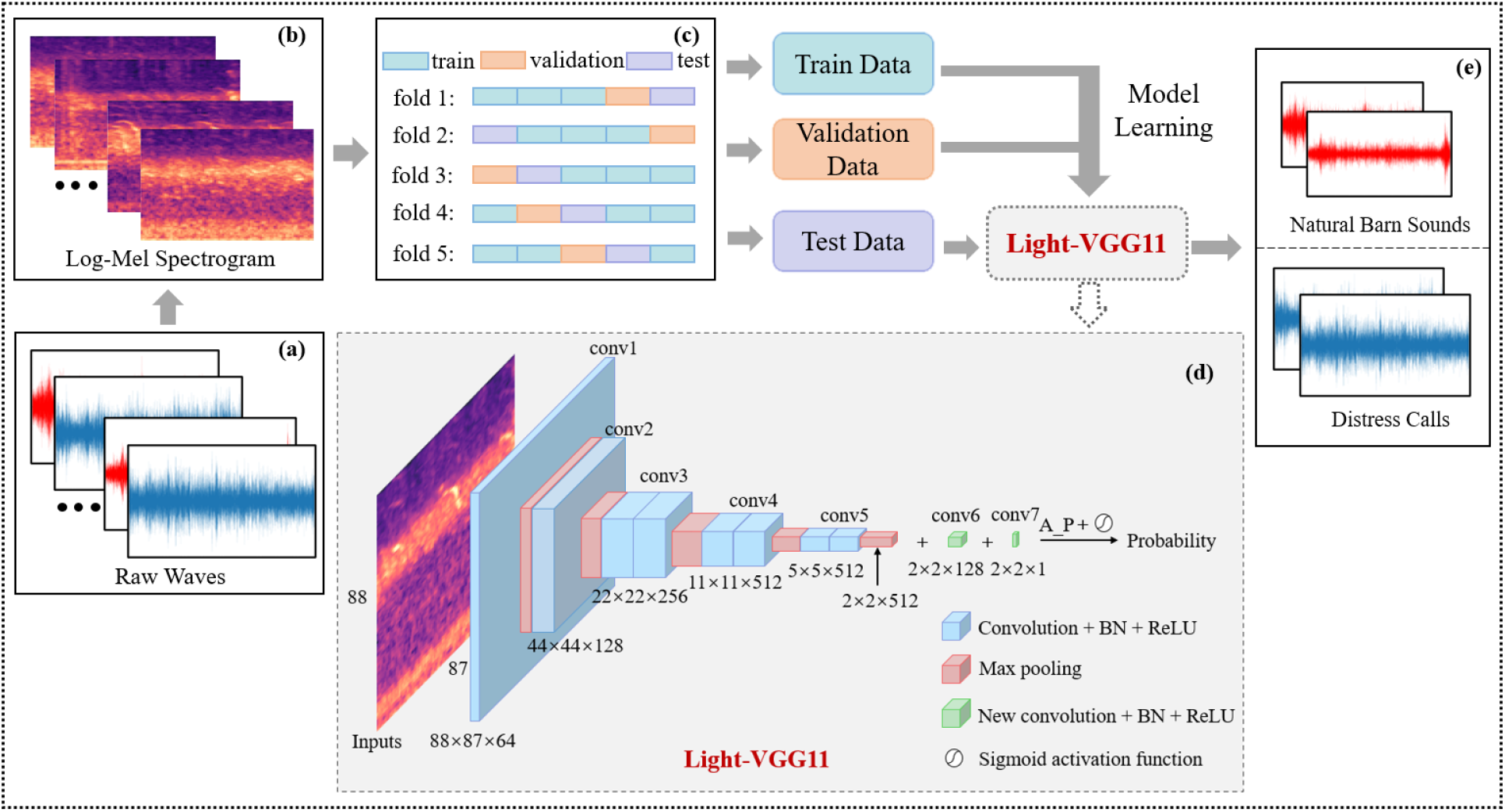
The overall flow chart of the audio classification method. (a) Raw waveform signals. (b) Log-Mel spectrograms. (c) Five-fold cross-validation. (d) The architecture of light-VGG11. (e) Predictions.

#### 2.3.1. Input Preparation

To represent audio signals as images within the use of CNNs, a log-Mel spectrogram, a typical and effective distinguishing feature tool in sound recognition [24,29,33,34], was applied for feature representation. We first applied STFT with Hann window function (length of 2,048 points) and 75% overlap (hop length of 512 points) to extract the magnitude spectrum [22,27,35], generating 87 temporal frames for each sample. Eighty-eight log-Mel scale band filters were then extracted rom the magnitude spectrum. Thus, a log-Mel spectrogram (88 × 87) was obtained as the two-dimensional image input, where the spectrum of frequencies (vertical y-axis, Hz) varied according to time (horizontal x-axis, sec) and the intensity of each pixel represented the amplitude (dB) of the bird call. The Librosa library [36] was used in this feature extraction process. The normalised spectrogram images are presented in Figure 4. We see that the high frequencies of natural barn sounds and distress calls mainly ranged from 2,048Hz to 4,096Hz, and distinction existed between these two sounds. Indeed, there are regular and repeated patterns occurring on distress calls as highlighted in the black boxes (Figure 4b) while we did not find any on natural barn sounds (Figure 4a).

**Figure 4.**
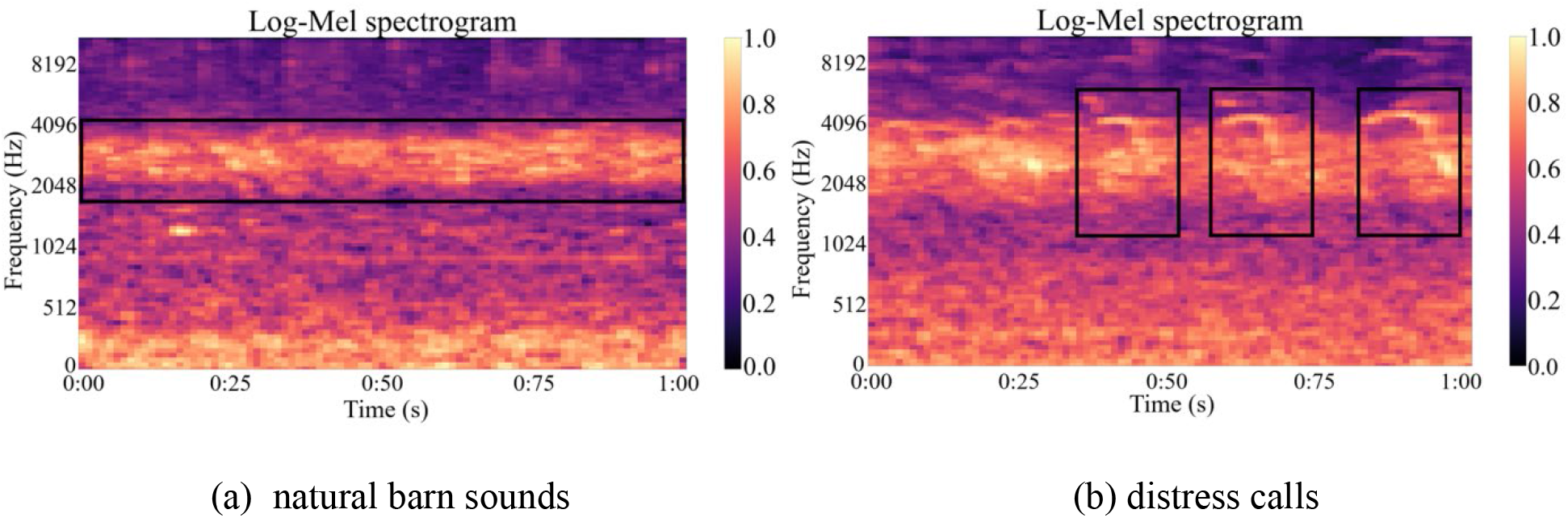
Log-Mel spectrograms samples of natural barn sounds and distress calls.

#### 2.3.2. Neural Network Structure

CNNs with very small (3 × 3) convolution filters, which effectively increase network depth and aggregate more complex patterns, have greatly succeeded in large-scale image classification [37]. This study adopted two types of popular CNNs, *i*.*e*., VGG Nets [37] and ResNets [38], to achieve the time-frequency representation learning. The preliminary results revealed that VGG11 displayed a favourable performance (Table 1). VGG11 network, which mainly consists of 11 weight layers, *i*.*e*., eight convolutional layers and three fully connected layers, has been widely used in various classification tasks [33,39]. However, these fully connected layers lead to the explosion of parameters (up to 128.8 million approximately), posing a challenge to the computation efficiency in real-world applications. Thus, we modified its structure to get a lightweight variant, renaming it as light-VGG11 (Figure 3d). Inspired by the architectures of ResNets [38], we removed the last three fully connected layers and replaced them with two convolutional layers followed by an average-pooling layer, effectively decreasing the number of parameters to 9.3 million. The operation was shown as follows:

**Table 1.** Comparative results between different models. The best results for each evaluation metric are highlighted in bold.

| Network<sup>#</sup> | Precision (%) | Recall (%) | F1 score (%) | Accuracy (%) | Para-meters (M) | Speed (ms) |
| --- | --- | --- | --- | --- | --- | --- |
| Machine learning |  |  |  |  |  |  |
| Naive Bayes | 72.01 | 73.40 | 72.14 | 73.03 | <1.0 | 6.74 |
| Decision Tree | 75.18 | 75.39 | 75.23 | 76.84 | <1.0 | 6.72 |
| SVM | 84.13 | 84.77 | 84.39 | 85.33 | <1.0 | 7.35 |
| Random Forest | 85.84 | 85.88 | 85.85 | 86.83 | <1.0 | 6.74 |
| GB | 87.40 | 87.64 | 87.51 | 88.33 | <1.0 | 6.73 |
| Deep learning |  |  |  |  |  |  |
| ResNet18 | 94.44 | 94.42 | 94.43 | 94.81 | 11.2 | 37.32 |
| ResNet34 | 94.33 | 94.42 | 94.38 | 94.75 | 21.3 | 75.96 |
| VGG11 | **95.16** | **95.43** | **95.29** | **95.60** | 128.8 | 84.66 |
| VGG16 | 94.79 | 95.13 | 94.95 | 95.28 | 134.3 | 129.75 |
| VGG19 | 94.91 | 95.24 | 95.07 | 95.39 | 139.6 | 151.52 |
| light-VGG11 | 94.58 | 94.89 | 94.73 | 95.07 | 9.3 | 54.31 |

<sup>#</sup>SVM: support vector machine; GB: gradient boosting.

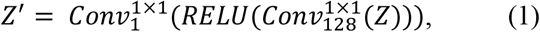

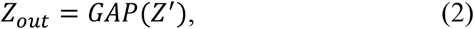

where *Z* and *Z*‘ denoted the feature maps after the last max-pooling layer and the last convolutional layer, respectively, and *Z*_*out*_ denoted the output. 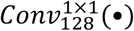 and 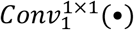 represent 1 × 1 convolution operations with 128 filters and one filter, respectively. *RELU*(•) denoted the rectified linear unit activation function [40], and *GAP*(•) denoted the global average-pooling operation. In addition, each convolution operation was followed by a batch normalisation operation. A sigmoid function was further applied to map the output to the probability between 0 and 1.

### 2.4. Data Augmentation

To artificially increase the diversity of the training dataset, inspired by previous research [30,41,42], we separately evaluated the impact of four different data augmentation strategies on the model’s performance, including time masking, frequency masking, mixed spectrograms of the same class (SpecSameClassMix), and Gaussian noise. Each method, when chosen, was randomly applied during training to each input image with a probability of 0.5. Figure 5 illustrates the visualisations of the spectrograms separately passed through these four augmentation methods. The details of these methods are shown as follows:

**Figure 5.**
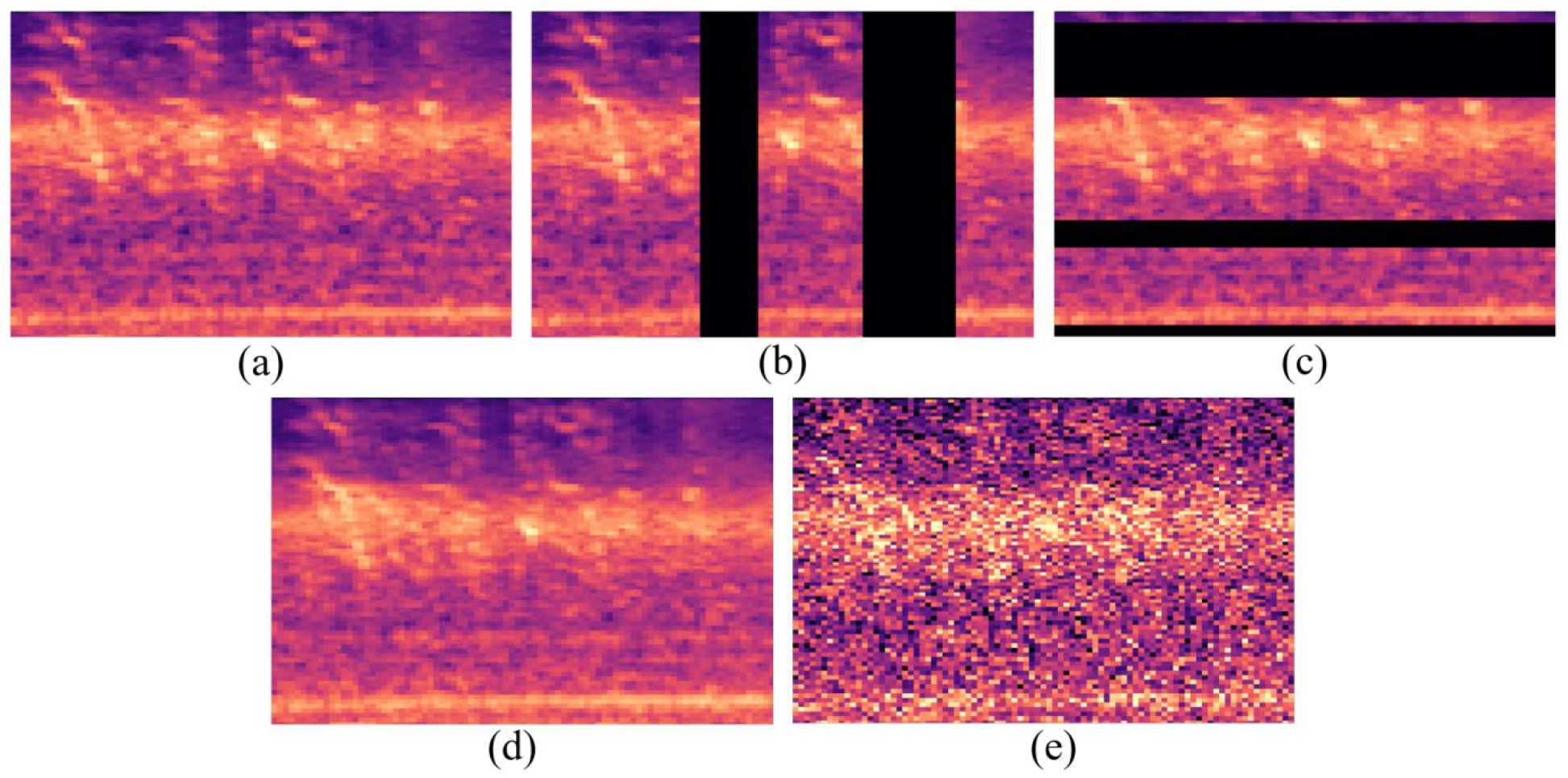
The original log-Mel spectrogram and its four transformed versions. (a) Original input. (b) Input with time masking. (c) Input with frequency masking. (d) Input with SpecSameClassMix. (e) Input with Gaussian noise.

1) Time masking. Time masking refers to the practice of randomly masking part of the spectrogram along with time-wise with a specific probability. It could effectively reduce the inter-dependence of features that appeared in the same image, thereby mitigating the information loss problem due to interruption of collected data caused by hardware devices. In our implementation, we adopted the time masking method proposed by Park *et al*. [30], *i*.*e*., two time masks with maximum adaptive size ratio (*p*_*s*_ =0.2), where the maximum adaptive size was set to *p*_*s*_ times the time dimensions of the spectrogram.

2) Frequency masking. Similar to time masking, we adopted the frequency masking method proposed by Park *et al*. [30] to enhance the model’s robustness, *i*.*e*., three frequency masks with parameter *F* = 15.

3) SpecSameClassMix. Inspired by the SpectrogramSameClassSum and Mixup methods [26,43], a new method was created here, namely SpecSameClassMix. Each mixed sample *z* was generated using *z* = *r* * *x* + (1 – *r*) * *y* where *x* was the original sample, *y* was the sample randomly selected from the same class, and *r* was chosen from a uniform distribution in the range [0, 1] [44].

4) Gaussian noise. Gaussian noise addition, a common data augmentation method, has been proven effective in some classification tasks [45,46]. The training data were augmented by adding noise generated from a Gaussian distribution with a mean of zero and a variance of σ^2^. Herein, σ was set as 0.3 [45].

### 2.5. Evaluation Metrics

The comprehensive performance of the audio classification model was indicated by the four commonly used evaluation metrics [24,47], *i*.*e*., precision, recall, F1score, and accuracy:

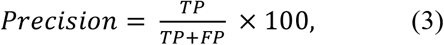

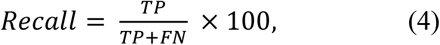

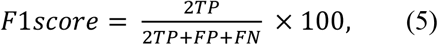

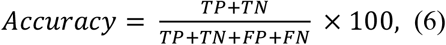

where *TP, FP, TN*, and *FN* were the number of true positives, false positives, true negatives, and false negatives, respectively. We calculated these four metrics of each category separately for the binary classification and then performed the macro average to eliminate the impacts of imbalance.

### 2.6. Implementation Details

To evaluate the effectiveness of deep learning models, we first tested five conventional machine learning methods, *i*.*e*., naïve Bayes (NB), decision tree (DT), SVM, random forest (RF), and gradient boosting (GB). During the feature extraction process, Mel Frequency Cepstrum Coefficients with 12 filter frequency bans were calculated based on the audio files of the distress calls and natural barn sounds [12]. These processed data were then scaled by removing the mean and scaling to unit variance, followed by being reshaped into one-dimensional arrays used for the final classification.

To obtain quick convergence and improve the model’s robustness and uncertainty, we pre-trained all selected deep learning models on the public dataset ImageNet and fine-tuned them using the current dataset. During training, the binary cross-entropy loss was chosen as the loss function, and an Adam optimiser with an initial learning rate of 1 × 10^−4^ was employed. The epochs and batch size were set to 100 and 128, respectively. The best model with the highest validation accuracy was saved and verified using test data. All experiments were executed using the PyTorch framework on an NVIDIA Tesla V100 GPU.

## 3. Results

### 3.1. Comparison of different models

We presented the comparative results of the light-VGG11 against machine learning methods (*i*.*e*., NB, DT, SVM, RF, and GB) and two kinds of deep learning methods (*i*.*e*., ResNets and VGG Nets) in Table 1. Note that we excluded the data augmentation processing here as we only attempted to evaluate the distinction of network architecture. As shown in Table 1, machine learning models obtained much lower values than deep learning algorithms in terms of all the metrics, although having significantly smaller parameter sizes and faster detection speed. It demonstrated the promising ability of deep learning for identifying chicken distress calls. Among the deep learning models, VGG Nets performed better than ResNets, where the VGG11 had the best performance with precision, recall, F1 score, and accuracy of 95.16%, 95.43%, 95.29, and 95.60%, respectively. However, the parameter sizes of VGGNets, which have reached more than one hundred million, are significantly larger than that of other networks. This required more computational costs and reduced operational efficiency, posing a challenge for online and real-time large data analysis in reality. In contrast, ResNet18 had the highest detection speed, as high as 37 ms, 126.85% faster than VGG11. Our light-VGG had 92.78% smaller parameters (9.3 million) and 55.88% faster detection speed than its previous architecture VGG11. In addition, it is worth noting that the light-VGG11 obtained a comparable performance with the precision, recall, F1 score, and accuracy of 94.58%, 94.89%, 94.73%, and 95.07%, respectively, in comparison with VGG11. It indicated that our light-VGG11 achieved a good trade-off between classification performance and computational cost.

### 3.2. Comparison of different models added with data augmentation

We implemented the network architectures of ResNet18, VGG11, and light-VGG11 and applied the same data augmentation method to the training set. The comparative results of these models are illustrated in Table 2. The light-VGG11, together with data augmentation, contributed to a favourable performance promotion, which demonstrated the good capability of data augmentation to enrich the limited training dataset in the current task. Specifically, the Gaussian noise possessed the best performance with precision, recall, F1 score, and accuracy of 95.40%, 95.81%, 95.60%, and 95.88%, respectively. Time masking enabled our model to obtain premium performance with precision, recall, F1 score, and accuracy of 95.49%, 95.11%, 95.30%, and 95.63%, respectively. Frequency masking and SpecSameClassMix also exhibited superior performance with different degrees of improvement, compared to not using data augmentation.

**Table 2.** Comparison between data augmentation strategies on ResNet18, VGG11, and light-VGG11. The best results for each metric are highlighted in bold.

| Network<sup>#</sup> | Data Aug | Precision (%) | Recall (%) | F1 score (%) | Accuracy (%) |
| --- | --- | --- | --- | --- | --- |
| ResNet18 | TM | 94.56 | 94.55 | 94.55 | 94.92 |
|  | FM | 94.52 | 94.46 | 94.50 | 94.87 |
|  | Spec_SCS | 92.29 | 93.51 | 93.88 | 94.34 |
|  | GN_A | 95.09 | 94.38 | 94.71 | 95.11 |
| VGG11 | TM | **95.72** | 95.13 | 95.41 | 95.75 |
|  | FM | 94.39 | 94.98 | 94.67 | 95.00 |
|  | Spec_SCS | 94.63 | 94.92 | 94.77 | 95.11 |
|  | GN_A | 95.44 | 95.48 | 95.46 | 95.77 |
| light-VGG11 | TM | 95.49 | 95.11 | 95.30 | 95.63 |
|  | FM | 95.25 | 94.79 | 95.01 | 95.37 |
|  | Spec_SCS | 94.88 | 94.82 | 94.85 | 95.20 |
|  | GN_A | 95.40 | **95.81** | **95.60** | **95.88** |

<sup>#</sup>No-Aug: without data augmentation; TM: time masking; FM: frequency masking; Spec\_SCS: SpecSameClassMix; GN\_A: Gaussian noise.

Our light-VGG11 always displayed a better performance than ResNet18, whichever data augmentation techniques were applied to them. In comparison to VGG11, our light-VGG11 showed superior capability for classification performance with increments of 0.86%, 0.34%, and 0.37% in precision, F1 score, and accuracy, respectively, when frequency masking was applied. We witnessed an increase of 0.25%, 0.08%, and 0.09% in precision, F1 score, and accuracy, respectively when SpecSameClassMix was applied. Finally, using Gaussian noise led to an increase of 0.33%, 0.20%, and 0.11% in the recall, F1 score, and accuracy, respectively. Hence, it can be concluded that the light-VGG11 outperformed ResNet18 and VGG11 when applied with data augmentation, which reinforced the suitability of our method to distress calls identification tasks using audio recordings in real scenarios especially in resource-constrained environments.

### 3.3. Identification performance analysis

Among these metrics, recall represents the percentage of correctly classified samples. The recall confusion matrix of our light-VGG11 without and with data augmentation is presented in Figure 6. We can see that the recall values of distress calls displayed improvements when data augmentation was applied, which revealed that data augmentation effectively enlarged the data representations of distress calls. Specifically, the time masking, frequency masking, SpecSameClassMix, and Gaussian noise enabled the recalls of distress calls to reach 97.12%, 97.03%, 96.28%, and 96.07%, respectively. In addition, it was worth noting that except for Gaussian noise which showed help in the detection of natural barn sounds, all other three data augmentation strategies degraded the recalls of natural barn sounds with different degrees. This indicated that more samples of barn sounds were misclassified as distress calls when applying time masking, frequency masking, as well as SpecSameClassMix.

**Figure 6.**
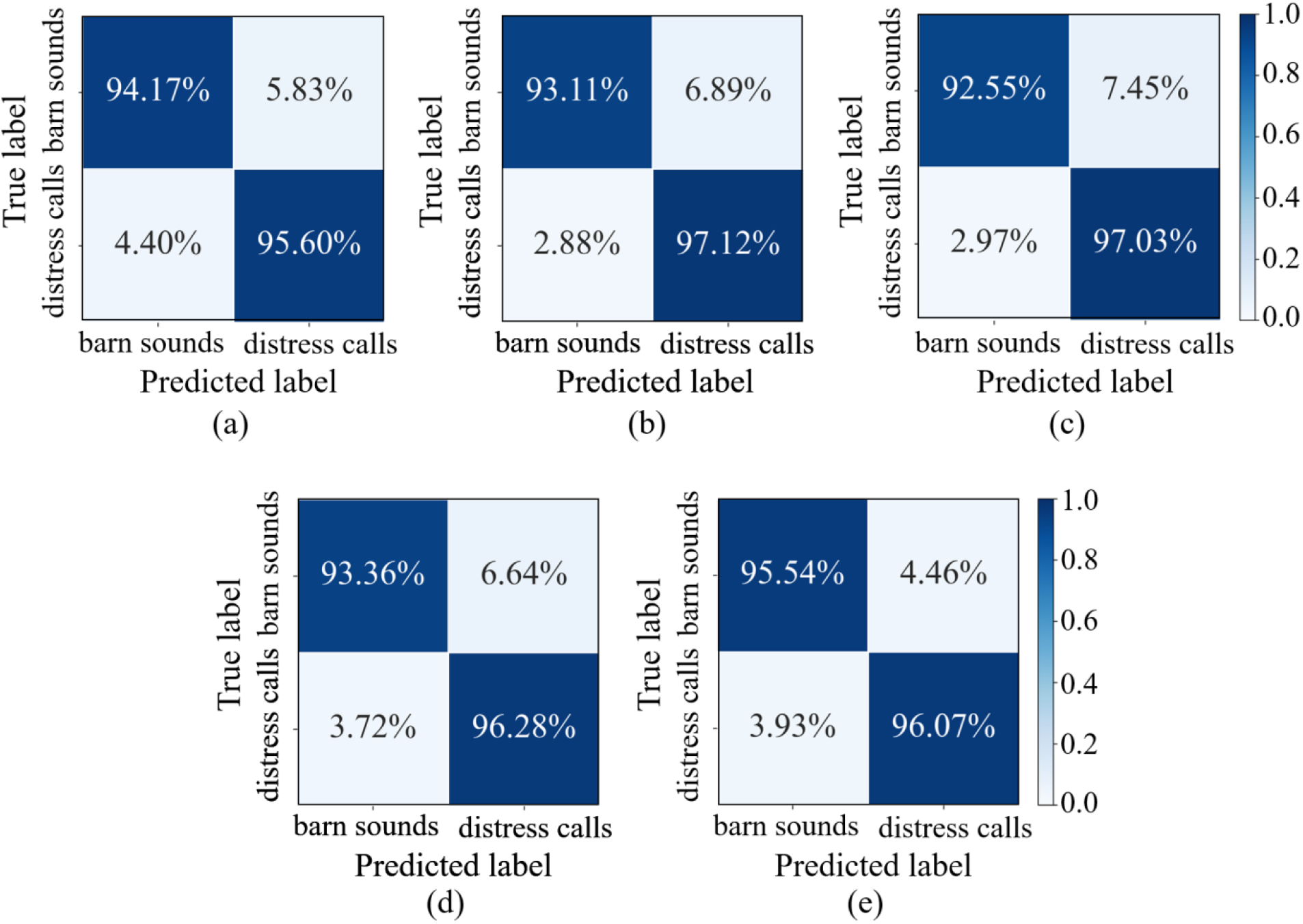
The recall confusion matrix of light-VGG11 without data augmentation (a) and with four data augmentation strategies: time masking (b), frequency masking (c), SpecSameClassMix (d), and Gaussian noise (e).

## 4. Discussion

In chickens, early-life welfare constraints often predict later life welfare concerns [48]. The output of distress vocalisations in commercial flocks is linked to growth rates and mortality levels [5]. However, since the annual global production of chickens exceeds 25 billion birds [49], and they are often housed in very large groups, numbering thousands, we need to develop automated methods and advanced technologies to monitor the production of distress calls. Herein we utilised deep learning classification methods to automatically identify chicken distress calls using audio recordings of chickens. Considering the practical implementation, we modified the structure of VGG11, which performed best preliminarily while possessing a large number of parameters, to get a novel light-VGG11 and trained it on the log-Mel spectrograms converted from raw audio signals. The results indicated that the light-VGG11 exhibited a comparable performance with 92.78% significantly smaller parameters and 55.88% faster detection speed than VGG11. Data augmentation was further adopted and validated to improve the classification performance of the light-VGG11 without increasing parameters. This illustrated that our method achieved distress calls identification at a low computational cost, giving it considerable potential for online and real-time large data analysis in real applications, particularly in scenarios with limited computation resources and power budgets [19,50].

To the best of our knowledge, our research is the first to exploit deep learning methods to identify chicken distress calls. It has shown its superior performance compared to conventional machine learning methods. This was because deep learning models could aggregate more complex and general patterns without domain knowledge. Among the selected deep learning models, the test performance of the ResNets was not as good as VGG Nets, although the shortcut connections in ResNets largely improved the ability of the model to learn some highly complex patterns in the data. This might be explained by the fact that the patterns in the current dataset were comparably not so complex to catch compared to the public dataset such as ImageNet [51]. The VGG11 outperformed other VGG Nets with deeper layers, further indicating that favourable performance can be obtained simply by using CNN models with shallower layers based on our dataset, as much deeper models tend to overfit. In addition, our light-VGG11 possessed competitive performance with VGG11 while having a significantly smaller parameter size and faster detection speed. This was because the convolutional layers in the raw architecture were kept, which effectively helps the network extract features.

Data augmentation was widely applied for improving the model’s generalisation ability in animal audio classification [26]. We found that data augmentation performed reasonably well on improving our model’s performance and particularly the detection rate of distress calls without increasing parameter size. This was ascribed to the fact that data augmentation took into account big amounts of noise or fluctuation presenting in real environments so as to better simulate real complexity [45]. In addition, all data augmentation strategies negatively impacted the detection of natural barn sounds except for the Gaussian noise technique which displayed better results. It reflected that Gaussian noise effectively promoted the robustness of the network to the environmental noise, which can be attributed to the fact that it is similar to the actual noise [52]. Beyond the data augmentation strategies that we used in our model, more complicated data augmentation methods are applied directly to audio recordings, such as time-stretching, pitch shifting, and mixing multiple audios [27,44,53]. However, those data augmentation techniques scale the dataset several times since it generates data-augmented samples before training models, taking up a large amount of storage space. On the contrary, the techniques used in our work were applied to spectrograms directly and performed on the fly during training only, which enabled the avoidance of storing the transformed images on the disk [28].

In the scenarios of real-world audio identification, it is necessary to consider the limited computational capabilities and constraints in resources, particularly low powered devices [42]. Some previous studies tended to reduce the computational cost by reducing model size and complexity [22,54], tuning the number of parameters [55], or replacing more efficient arithmetic operations [56]. Anvarjon *et al*. [57] applied a lightweight CNN model with fewer layers for speech emotion recognition. A lightweight human action recognition algorithm based on one-dimensional CNN was verified to outperform previous solutions in energy efficiency, with much fewer parameters (1.3 million) instead of more than 88 million parameters of commonly used networks [50]. In this study, our light-VGG11 was more lightweight and competent for big data analysis as well as real-time monitoring since it improved the primitive VGG11 to achieve identification of distress calls at a low computational cost. In future, this method will potentially allow staff to monitor chicken welfare in real-time and remotely, promoting earlier husbandry interventions when necessary. It can also reduce analysts’ workloads and facilitate the analysis of large datasets, improving the husbandry and management of animals [22].

We investigated one type of vocalisation (*i*.*e*., distress call), and the recordings came from one particular field season. In real-world identification, many other specific chicken sounds like alarm and gakel calls may also be regarded as potential welfare indicators [17], and distress calls might be inconsistent between different breeds and welfare scale systems. This will require us to build a larger dataset that incorporates more types of vocalisations from different breeds and production environments in the future. In addition, our model could be integrated into more complete systems with other detection methods to achieve additional functions like identifying the vocal source location [58].

Overall, our method showed a potential possibility for identifying chicken distress calls using deep learning combined with bioacoustic techniques, allowing the development of technologies that can monitor the output of distress calls in large, commercial chicken flocks. It also fully considered the constraints in computation resources by slightly modifying the existing CNN architecture to reduce time overhead, and the difficulty of acquiring real-world datasets by utilising data augmentation to increase dataset diversity. As part of a precision livestock farming system, it is crucial to interpret how the outputs of distress calls reflect their external surroundings [59,60]. Thus, future work is also required to construct an integrated system with the potential to grasp the environmental condition of chickens through the analysis of audio recordings, ultimately contributing to chicken welfare.

## Ethics

This study was reviewed and approved by the Animal Welfare and Ethical Review Board committee of Queen Mary University of London.

## Data accessibility

Data for the analyses reported can be accessed via https://drive.google.com/file/d/1OlQdWnAUxaeq6n2s6tWSz68u-GLIWloZ/view?usp=sharing. The source code is available at https://github.com/Max-1234-hub/light-VGG11.

## Authors’ contributions

The study was conceptualized by Z.Q.X., Z.X.X, Y.G., J.T., D.B., R.S., and A.G.M. The data were collected by Z.Q.X., and Z.X.X. Data analyses were conducted by A.X.M, K.L., and I.D.A.N. The manuscript was drafted by A.X.M., C.G., K.L., R.S., and A.G.M. All authors approved the final version of the manuscript and agree to be held accountable for the content of this paper.

## Acknowledgments

We thank Michael Mcloughlin and Ben McCarthy for their help, and the farmers for access to their animals.

## References

1. Briefer EF, Tettamanti F, McElligott AG. 2015 Emotions in goats: mapping physiological, behavioural and vocal profiles. Anim. Behav. 99, 131–143. (doi:10.1016/j.anbehav.2014.11.002)

2. Friel M, Kunc HP, Griffin K, Asher L, Collins LM. 2019 Positive and negative contexts predict duration of pig vocalisations. Sci. Rep. 9, 1–7. (doi:10.1038/s41598-019-38514-w)

3. McGrath N, Dunlop R, Dwyer CD, Burman O, Phillips CJC. 2017 Hens vary their vocal repertoire and structure when anticipating different types of reward. Anim. Behav. 130, 79–96. (doi:10.1016/j.anbehav.2017.05.025)

4. Warnick JE, Huang CJ, Acevedo EO, Sufka KJ. 2009 Modelling the anxiety-depression continuum in chicks. J. Psychopharmacol. 23, 143–156. (doi:10.1177/0269881108089805)

5. Herborn KA, McElligott AG, Mitchell MA, Sandilands V, Bradshaw B, Asher L. 2020 Spectral entropy of early-life distress calls as an iceberg indicator of chicken welfare. J. R. Soc. Interface 17. (doi:10.1098/rsif.2020.0086)

6. Manteuffel G, Puppe B, Schön PC. 2004 Vocalization of farm animals as a measure of welfare. Appl. Anim. Behav. Sci. 88, 163–182. (doi:10.1016/j.applanim.2004.02.012)

7. Carroll BT, Anderson DV, Daley W, Harbert S, Britton DF, Jackwood MW. 2014 Detecting symptoms of diseases in poultry through audio signal processing. In 2014 IEEE Global Conference on Signal and Information Processing (GlobalSIP), pp. 1132–1135. Atlanta, USA. (doi:10.1109/GlobalSIP.2014.7032298)

8. Fontana I, Tullo E, Butterworth A, Guarino M. 2015 An innovative approach to predict the growth in intensive poultry farming. Comput. Electron. Agric. 119, 178–183. (doi:10.1016/j.compag.2015.10.001)

9. Rizwan M, Carroll BT, Anderson DV, Daley W, Harbert S, Britton DF, Jackwood MW. 2016 Identifying rale sounds in chickens using audio signals for early disease detection in poultry. In 2016 IEEE Global Conference on Signal and Information Processing (GlobalSIP), pp. 55–59. Greater Washington, USA. (doi:10.1109/GlobalSIP.2016.7905802)

10. Whitaker BM, Carroll BT, Daley W, Anderson DV. 2014 Sparse decomposition of audio spectrograms for automated disease detection in chickens. In 2014 IEEE Global Conference on Signal and Information Processing (GlobalSIP), pp. 1122–1126. Atlanta, USA. (doi:10.1109/GlobalSIP.2014.7032296)

11. Favaro L, Briefer EF, McElligott AG. 2014 Artificial Neural Network Approach for Revealing Individuality, Group Membership and Age Information in Goat Kid Contact Calls. Acta Acust. United Acust. 100, 782–789. (doi:10.3813/AAA.918758)

12. Mcloughlin MP, Stewart R, McElligott AG. 2019 Automated bioacoustics: methods in ecology and conservation and their potential for animal welfare monitoring. J. R. Soc. Interface 16. (doi:10.1098/rsif.2019.0225)

13. Stowell D. 2018 Computational Bioacoustic Scene Analysis. In Computational Analysis of Sound Scenes and Events (eds T Virtanen, MD Plumbley, D Ellis), pp. 303–333. Cham: Springer International Publishing. (doi:10.1007/978-3-319-63450-0_11)

14. Stowell D, Petrusková T, Šálek M, Linhart P. 2019 Automatic acoustic identification of individuals in multiple species: improving identification across recording conditions. J. R. Soc. Interface 16. (doi:10.1098/rsif.2018.0940)

15. Stowell D, Wood MD, Pamuła H, Stylianou Y, Glotin H. 2019 Automatic acoustic detection of birds through deep learning: The first Bird Audio Detection challenge. Methods Ecol. Evol. 10, 368–380. (doi:10.1111/2041-210X.13103)

16. Carpentier L, Vranken E, Berckmans D, Paeshuyse J, Norton T. 2019 Development of sound-based poultry health monitoring tool for automated sneeze detection. Comput. Electron. Agric. 162, 573–581. (doi:10.1016/j.compag.2019.05.013)

17. Du X, Carpentier L, Teng G, Liu M, Wang C, Norton T. 2020 Assessment of Laying Hens’ Thermal Comfort Using Sound Technology. Sensors 20, 1–14. (doi:10.3390/s20020473)

18. Mao A, Huang E, Gan H, Parkes RSV, Xu W, Liu K. 2021 Cross-Modality Interaction Network for Equine Activity Recognition Using Imbalanced Multi-Modal Data. Sensors 21, 5818. (doi:10.3390/s21175818)

19. Du X, Teng G, Wang C, Carpentier L, Norton T. 2021 A tristimulus-formant model for automatic recognition of call types of laying hens. Comput. Electron. Agric. 187, 106221. (doi:10.1016/j.compag.2021.106221)

20. Dokuz Y, Tufekci Z. 2021 Mini-batch sample selection strategies for deep learning based speech recognition. Appl. Acoust. 171. (doi:10.1016/j.apacoust.2020.107573)

21. Jung D-H, Kim NY, Moon SH, Kim HS, Lee TS, Yang J-S, Lee JY, Han X, Park SH. 2021 Classification of Vocalization Recordings of Laying Hens and Cattle Using Convolutional Neural Network Models. J. Biosyst. Eng. 46, 217–224. (doi:10.1007/s42853-021-00101-1)

22. Madhusudhana S et al. 2021 Improve automatic detection of animal call sequences with temporal context. J. R. Soc. Interface 18. (doi:10.1098/rsif.2021.0297)

23. Badshah A, Ahmad J, Rahim N, Baik S. 2017 Speech Emotion Recognition from Spectrograms with Deep Convolutional Neural Network. In 2017 International Conference on Platform Technology and Service (PlatCon), pp. 1–5. Busan, Korea. (doi:10.1109/PlatCon.2017.7883728)

24. Nasirahmadi A, Gonzalez J, Sturm B, Hensel O, Knierim U. 2020 Pecking activity detection in group-housed turkeys using acoustic data and a deep learning technique. Biosyst. Eng. 194, 40–48. (doi:10.1016/j.biosystemseng.2020.03.015)

25. Zhong M, Taylor R, Bates N, Christey D, Basnet H, Flippin J, Palkovitz S, Dodhia R, Lavista Ferres J. 2021 Acoustic detection of regionally rare bird species through deep convolutional neural networks. Ecol. Inform. 64. (doi:10.1016/j.ecoinf.2021.101333)

26. Nanni L, Maguolo G, Paci M. 2020 Data augmentation approaches for improving animal audio classification. Ecol. Inform. 57, 1–26. (doi:10.1016/j.ecoinf.2020.101084)

27. Greco A, Petkov N, Saggese A, Vento M. 2020 AReN: A Deep Learning Approach for Sound Event Recognition Using a Brain Inspired Representation. IEEE Trans. Inf. Forensics 15, 3610–3624. (doi:10.1109/TIFS.2020.2994740)

28. Pellegrini T. 2017 Densely Connected CNNs for Bird Audio Detection. In 25th European Signal Processing Conference (EUSIPCO), pp. 1734–1738. Kos, Greece. (doi:10.23919/EUSIPCO.2017.8081506)

29. Park DS, Chan W, Zhang Y, Chiu C-C, Zoph B, Cubuk ED, L. QV. 2019 SpecAugment: A Simple Data Augmentation Method for Automatic Speech Recognition. In Proceedings of the Annual Conference of the International Speech Communication Association, INTERSPEECH, pp. 2613–2617. Graz, Austria. (doi:10.21437/Interspeech.2019-2680)

30. Park DS, Zhang Y, Chiu C, Chen Y, Li B, Chan W, Le QV, Wu Y. 2020 Specaugment on Large Scale Datasets. In ICASSP, IEEE International Conference on Acoustics, Speech and Signal Processing, pp. 6879–6883. Barcelona, Spain. (doi:10.1109/ICASSP40776.2020.9053205)

31. Cannam C, Landone C, Sandler M. 2010 Sonic visualiser: an open source application for viewing, analysing, and annotating music audio files. In Proceedings of the 18th ACM international conference on Multimedia, pp. 1467–1468. New York, USA: Association for Computing Machinery. (doi:10.1145/1873951.1874248)

32. Collias NE. 1987 The Vocal Repertoire of the Red Junglefowl: A Spectrographic Classification and the Code of Communication. The Condor 89, 510–524. (doi:10.2307/1368641)

33. Liu X, Iqbal T, Zhao J, Huang Q, Plumbley MD, Wang W. 2021 Conditional Sound Generation Using Neural Discrete Time-Frequency Representation Learning. In IEEE 31st International Workshop on Machine Learning for Signal Processing (MLSP), pp. 25–28. Gold Coast, Australia.

34. Zhao J, Mao X, Chen L. 2019 Speech emotion recognition using deep 1D & 2D CNN LSTM networks. Biomed. Signal Process. Control 47, 312–323. (doi:10.1016/j.bspc.2018.08.035)

35. Sprengel E, Jaggi M, Kilcher Y, Hofmann T. 2016 Audio Based Bird Species Identification using Deep Learning Techniques. In Conference and Labs of the Evaluation Forum, pp. 547–559. Evoria, Portugal.

36. McFee B, Raffel C, Liang D, Ellis D, McVicar M, Battenberg E, Nieto O. 2015 librosa: Audio and Music Signal Analysis in Python. In Python in Science Conference, pp. 18–24. Austin, Texas. (doi:10.25080/Majora-7b98e3ed-003)

37. Simonyan K, Zisserman A. 2015 Very Deep Convolutional Networks for Large-Scale Image Recognition. In 3rd International Conference on Learning Representations, pp. 1–14. San Diego, USA.

38. He K, Zhang X, Ren S, Sun J. 2016 Deep Residual Learning for Image Recognition. In 2016 IEEE Conference on Computer Vision and Pattern Recognition (CVPR), pp. 770–778. Las Vegas, USA. (doi:10.1109/CVPR.2016.90)

39. Duan C, Yin P, Zhi Y, Li X. 2019 Image Classification of Fashion-mnist Data Set Based on VGG Network. In 2019 2nd International Conference on Information Science and Electronic Technology, pp. 18–21. Taiyuan, China: Clausius Scientific Press. (doi:10.23977/iset.2019.004)

40. Nair V, Hinton GE. 2010 Rectified linear units improve restricted boltzmann machines. In Proceedings of the 27th International Conference on International Conference on Machine Learning, pp. 807–814. Madison, USA. (doi:10.1123/jab.2016-0355)

41. Nanni L, Costa YMG, Aguiar RL, Mangolin RB, Brahnam S, Silla CN. 2020 Ensemble of convolutional neural networks to improve animal audio classification. EURASIP Journal on Audio, Speech, and Music Processing 2020, 8. (doi:10.1186/s13636-020-00175-3)

42. Han W, Zhang Z, Zhang Y, Yu J, Chiu C-C, Qin J, Gulati A, Pang R, Wu Y. 2020 ContextNet: Improving Convolutional Neural Networks for Automatic Speech Recognition with Global Context. In Proceedings of the Annual Conference of the International Speech Communication Association, INTERSPEECH, pp. 3610–3614. Shanghai, China. (doi:10.21437/Interspeech.2020-2059)

43. Zhang H, Cisse M, Dauphin YN, Lopez-Paz D. 2018 mixup: Beyond Empirical Risk Minimization. In ICLR 2018 Conference, pp. 1–13. Vancouver, Canada.

44. Salamon J, Bello JP. 2017 Deep Convolutional Neural Networks and Data Augmentation for Environmental Sound Classification. In IEEE Signal Processing Letters, pp. 279–283. (doi:10.1109/LSP.2017.2657381)

45. Eerdekens A, Deruyck M, Fontaine J, Martens L, Poorter ED, Plets D, Joseph W. 2020 Resampling and Data Augmentation For Equines’ Behaviour Classification Based on Wearable Sensor Accelerometer Data Using a Convolutional Neural Network. In 2020 International Conference on Omni-layer Intelligent Systems (COINS), pp. 1–6. Barcelona, Spain. (doi:10.1109/COINS49042.2020.9191639)

46. Haradal S, Hayashi H, Uchida S. 2018 Biosignal Data Augmentation Based on Generative Adversarial Networks. In 2018 40th Annual International Conference of the IEEE Engineering in Medicine and Biology Society (EMBC), pp. 368–371. Honolulu, USA. (doi:10.1109/EMBC.2018.8512396)

47. Duan G, Zhang S, Lu M, Okinda C, Shen M, Norton T. 2021 Short-term feeding behaviour sound classification method for sheep using LSTM networks. Int. J. Agric. Biol. 14, 43–54. (doi:10.25165/ijabe.v14i2.6081)

48. Rodenburg TB, Komen H, Ellen ED, Uitdehaag KA, van Arendonk JAM. 2008 Selection method and early-life history affect behavioural development, feather pecking and cannibalism in laying hens: A review. Appl. Anim. Behav. Sci. 110, 217–228. (doi:10.1016/j.applanim.2007.09.009)

49. Shahbandeh M. 2021 Number of chickens worldwide from 1990 to 2019 (in million animals). statista. See https://www.statista.com/statistics/263962/number-of-chickens-worldwide-since-1990/#statisticContainer. (accessed on 1 December 2021).

50. Zhang B, Han J, Huang Z, Yang J, Zeng X. 2019 A Real-Time and Hardware-Efficient Processor for Skeleton-Based Action Recognition With Lightweight Convolutional Neural Network. IEEE Trans. Circuits Syst. II: Express Br. 66, 2052–2056. (doi:10.1109/TCSII.2019.2899829)

51. Wang Z, Yan W, Oates T. 2016 Time Series Classification from Scratch with Deep Neural Networks: A Strong Baseline. In Proceedings of the International Joint Conference on Neural Networks, pp. 1578–1585. Vancouver, Canada. (doi:10.1109/IJCNN.2017.7966039)

52. Kim E, Kim J, Lee H, Kim S. 2021 Adaptive Data Augmentation to Achieve Noise Robustness and Overcome Data Deficiency for Deep Learning. Appl. Sci. 11, 5586. (doi:10.3390/app11125586)

53. Lostanlen V, Salamon J, Farnsworth A, Kelling S, Bello JP. 2019 Robust sound event detection in bioacoustic sensor networks. PLoS One 14, e0214168. (doi:10.1371/journal.pone.0214168)

54. Sun H, Xu H, Liu B, He D, He J, Zhang H, Geng N. 2021 MEAN-SSD: A novel real-time detector for apple leaf diseases using improved light-weight convolutional neural networks. Comput. Electron. Agric. 189. (doi:10.1016/j.compag.2021.106379)

55. Eerdekens A, Deruyck M, Fontaine J, Martens L, De Poorter E, Plets D, Joseph W. 2021 A framework for energy-efficient equine activity recognition with leg accelerometers. Comput. Electron. Agric. 183, 106020. (doi:10.1016/j.compag.2021.106020)

56. Edel M, Köppe E. 2016 Binarized-BLSTM-RNN based Human Activity Recognition. In 2016 International Conference on Indoor Positioning and Indoor Navigation (IPIN), pp. 1–7. Sapporo, Japan. (doi:10.1109/IPIN.2016.7743581)

57. Anvarjon T, Mustaqeem Kwon S. 2020 Deep-Net: A Lightweight CNN-Based Speech Emotion Recognition System Using Deep Frequency Features. Sensors 20, 1–16. (doi:10.3390/s20185212)

58. Silva M, Ferrari S, Costa A, Aerts J-M, Guarino M, Berckmans D. 2008 Cough localization for the detection of respiratory diseases in pig houses. Comput. Electron. Agric. 64, 286–292. (doi:10.1016/j.compag.2008.05.024)

59. Green AC, Johnston IN, Clark CEF. 2018 Invited review: The evolution of cattle bioacoustics and application for advanced dairy systems. animal 12, 1250–1259. (doi:10.1017/S1751731117002646)

60. Stachowicz J, Umstätter C. 2021 Do we automatically detect health-or general welfare-related issues? A framework. Proc. Royal Soc. B 288. (doi:10.1098/rspb.2021.0190)

